# Modeling glioma-induced impairments on the glymphatic system

**DOI:** 10.1101/2025.04.16.648938

**Authors:** Alexandre Poulain, Jørgen Riseth, Kyrre E. Emblem, Kent-Andre Mardal

**Author notes:** These authors contributed equally to this work.

## Abstract

Altered glymphatic function is observed for many neurological diseases. Glioma, one of the most common brain cancers, is known to have altered fluid dynamics in terms of edema and blood-brain-barrier breakdown, both features potentially impacting the glymphatic function. To study glioma and its fluid dynamics, we propose a flexible mathematical model, including the tumor, the peri-tumoral edema as well as the healthy tissue. From a mechanical point of view, we consider the brain as a multicompartment porous medium and model both the fluid movement in the brain and the clearance of solutes that are convected but also diffuse in the extra-cellular space. Our results indicate that the impairment on the glymphatic system due to glioma growth is two-fold. First, edema resulting from leakage of fluid from the blood vessels or occlusion of the peri-vascular spaces, considered as an exit route, result in a local high pressure zone, consequently impairing negatively glymphatic clearance. Second, local changes of porosities (volume fraction of certain compartments such as perivascular or extracellular spaces), result in a disruption of the transport of solutes in the brain. Our results indicate that an effect similar to the enhanced permeability and retention is obtained using biologically relevant changes of parameter values of our model. Our mathematical model can be viewed as the first step towards a digital twin for drug or contrast product delivery within the cerebro-spinal fluid directly (*e*.*g*. from intrathecal injection) for patients suffering from gliomas.

## 1 Introduction

Cancers of the brain and the central nervous system (CNS) have a relatively low incidence, but remain among the deadliest cancers [22]. One of the most common type is glioma and one of its features is an altered fluid dynamics system. In particular, gliomas are typically associated with a peritumoral edema [48, 38, 47], which is clearly visible using various Magnetic Resonance (MR) imaging techniques [54].

The understanding of the fluid dynamics of the brain has changed dramatically the last decade. In particular, the glymphatic system proposes an alternative view on the extra-vascular fluid dynamics, where cerebrospinal fluid filled perivascular spaces are tightly coupled to the interstitial fluid in a brain-wide network that facilitates long range solute transfer and clearance [30]. While the mechanisms behind the glymphatic system is still actively investigated, and multiple driving forces has been proposed [8, 23, 45], it is clear from numerical modeling that it amplifies solute transfer in both rodents [51] and humans [65] as compared to extra-cellular diffusion alone. To what extent the glymphatic system is a major component of gliomas and their development is currently unknown.

Tumors and associated edema are known to have altered fluid dynamics as compared to healthy tissue. The apparent diffusion coefficient (ADC) of a tumor is very heterogeneous compared to that of healthy tissue, whereas edema regions tend to have higher ADC. Further, the edema is associated with alterations both in porosity and permeability in or between some of the brain and vasculature structures involved in the glymphatic system, depending on the type of edema. First, vasogenic edema is characterized by a breakdown of the blood-brain-barrier, and as such an increased permeability between the vasculature and the associated perivascular space (PVS) [15]. Second, non-vasogenic edema is associated with increased cellular volume either causing or caused by osmotic gradients [37]. To the authors knowledge, only the work of Siri *et al*. [55] dealt with the modeling of the the effect of glioma and its associated edema in the context of glymphatics. In this work, the authors proposed to analyze using a simple and tractable mechanical model and an electrical analog model the effect of the size and location of the tumor on compression and decreased fluid flow in the PVS. However, their models do not represent pressure and fluid flow at the scale of the brain nor the evolution of the distribution of solutes within the brain. As such, we here propose a multi-compartment model for the solute transfer and interactions between the tumor, edema and surrounding healthy brain tissue.

Intrathecal injection of Gadobutrol [52, 17, 18], in contrast to an intravenous injection, puts the focus on extra-vascular pathways and barriers of the CNS as Gadobutrol do not cross the blood-brain barrier to a significant extent. Intrathecal injection is recognized as having a significant potential for drug-delivery [6, 10], but the modeling is challenging in particular because of the complex CSF flow [27, 3, 32] that varies significantly between individuals [16]. We consider the distribution within the brain in a modeling framework similar to [62, 65, 14], where simulation results, given boundary conditions provided by imaging, yield good approximations of the glymphatic transport.

Here, we propose a model for investigating the solute transfer of the glymphatic system in patients with gliomas. We propose a six compartment system where the solute transfer in and between the vascular, perivascular, and extra-cellular compartment are modeled via compartment pressures and solute concentrations. The model was previously proposed and verified in a murine model [49], although gliomas where not considered. Here, we extend the model with a glioma and its associated peri-tumor edema, modeled in terms of altered porosity and permeability. We use a finite element model of a complete human brain taken from [42] to enable brain-wide simulations of both different models of gliomas and peri-tumor environments.

## 2 Methods

### 2.1 Mathematical notations

#### Domains

We denote Ω ⊂ ℝ^3^ the brain domain, and assume that its boundary *∂*Ω_*B*_ ∪ *∂*Ω_ventr_ of this domain is sufficiently smooth (*∂*Ω_*B*_ denotes the surface of the brain in contact with the pia mater and *∂*Ω_ventr_ denotes the surface lining the walls of the ventricles, *i*.*e*. where the pia mater joins the ependyma to form the choroid plexus). Let **x** ∈ Ω be any point of this domain and its coordinates be given by **x** = (*x*_1_, *x*_2_, *x*_3_). In this article, bold symbols are used to denote vectors. The time-space domain is denoted by Ω_*T*_ = Ω × (0, *T*) for some finite time horizon *T >* 0. Similarly, we use the notation *∂*Ω_*T*_ = *∂*Ω × (0, *T*). The brain tissue is modeled as different separated compartments described below.

#### Compartments within the brain

We model the healthy brain tissue as consisting of four structures, namely blood vessels, perivascular spaces, extracellular space and the solid skeleton formed by the cells. We further subdivide the blood vessels and the perivascular spaces into (peri-)arterial and (peri-)venous compartments. In total, this constitutes 6 compartments:

- Cells/intracellular space *s*.
- Extracellular space *e*.
- Perivascular spaces 𝒫= {*pa, pv*} around the arteries (and arterioles) and veins (and venules).
- Blood vessels ℬ= {*a, v*} consisting of arteries, arterioles, veins and venules.

### 2.2 Modeling assumptions

As for every mathematical model, we make the simplifying assumptions to reduce the complexity of the problem and derive a tractable mathematical model. We use the following assumptions:

**(A1)** The tissue is rigid on the relevant time scale.

**(A2)** Exchange of both fluids and solutes between the cell interior and their surroundings is negligible (*c*_*s*_ = 0, *p*_*s*_ = constant), and is therefore excluded from the model.

**(A3)** Clearance of tracer that entered the blood is faster than the time scale of interest (*c*_*j*_ = 0, *j* ∈ℬ),

**(A4)** No direct pathway from ECS to blood (tracers and fluid must pass through the PVS compartment).

**(A5)** For an intact BBB, we assume there is no exchange of fluid or solute between PVS and blood compartments.

**(A6)** We do not model explicitly the dynamics of the fluid in the blood compartments. We assume a constant pressure field *p*_*j*_ for *j* ∈ B.

### 2.3 Mathematical modelling of the glymphatic system

We propose a multicompartment model of the glymphatic system to represent both fluid movement and transport of solutes within the parenchyma. Our model can be derived from a formal volume averaging method (see Appendix A). Following Assumptions (A1)–(A6) as well as the fact that inertia effects are negligible and CSF is incompressible, we obtain a static description of fluid flow (*i*.*e*. meaning that the equations for the fluid do not depend explicitly on time), while solute transport is dynamic in time (*i*.*e*. time-dependent). As a consequence, velocity fields **v**_**j**_(*x*) and pressures *p*_*j*_(*x*) depend solely on space whereas concentrations *c*_*j*_(*t, x*) depend both on time and space. In the following, we will generally drop the indication of the dependencies on time and space.

#### Fluid flow in the brain parenchyma and PVSs

The brain tissue is considered as a porous medium. Fluid velocity is computed using Darcy’s law, *i*.*e*. velocity is given by the opposite direction of the fluid pressure gradient times the permeability of the compartment, or stated mathematically

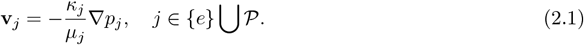

In the previous equation, *κ*_*j*_ denotes the hydraulic permeability of the tissue within the compartment, and *µ*_*j*_ is the dynamic viscosity of the fluid (interstitial fluid or CSF).

CSF and interstitial fluid flows in the brain are modeled using a multicompartment porous medium (see *e*.*g*. [49]). For *j* ∈ {*e*} ∪ 𝒫, we have

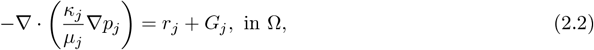

with *r*_*j*_ a transfer function that represents transfer of fluid between the compartments and *G*_*j*_ a possible source or sink of fluid (this latter term will be used to represent potential leakage from blood vessels due to disrupted BBB).

#### Pressure at the pial surface and boundary conditions

The pressure boundary conditions are given by Robin type boundary conditions for ∈ *j* {*e*} ⋃ 𝒫to represent fluid transmission at the pial surface, *i*.*e*.

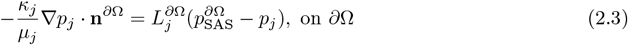

where 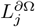 denotes the hydraulic conductivity at the pial surface, 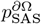 denotes the fluid pressure applied at the pial surface from CSF in SAS (assumed to be constant on *∂*Ω), and **n**^*∂*Ω^ is the outward normal vector to the boundary *∂*Ω. Note that 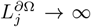 correspond to a Dirichlet boundary condition with 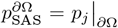(where *p*_*j* *∂*Ω_ is the restriction of *p*_*j*_ to the boundary *∂*Ω), whereas 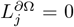 corresponds to a homogeneous Neumann boundary condition for the pressure.

#### Fluid transfer between the compartments

The function *r*_*j*_ represents the transfer of fluid between the compartment *j* ∈ {*e*} ⋃ 𝒫 and the other compartments, *i*.*e*.

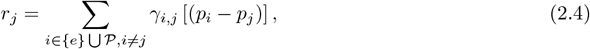

where *γ*_*i,j*_ represents the permeability and the surface to volume ratio of the membrane separating compartments *i* and *j, i*.*e*.

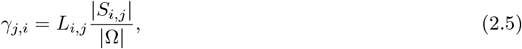

where |Ω| = 𝒫_Ω_ 1 d**x** denotes the volume of the brain, and *L*_*i,j*_ is the hydraulic conductivity of the membrane separating the *i*−th and *j*−th compartments, with 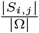 being the ratio between the surface of the membrane and the volume of the tissue.

#### Fluid leakage from the blood compartments to represent leaky blood vessels

To represent the leakage of fluid from the blood vessels that could occur in a vasogenic edema, we use

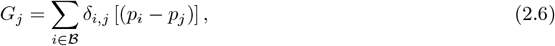

with the permeability of the discrupted vessels given by *δ*_*i,j*_ = 𝟙_Tumor_*α*_*i,j*_*γ*_*i,j*_, where 1_Tumor_ is the characteristic function to localize this effect in the tumor region and *α*_*i,j*_ *>* 1 a prefactor that represents the increase of the blood vessel permeability.

#### Glymphatic solute transport in the brain

We model solute transport within the brain following the same approach as in [49]. For all **x** ∈ Ω, *t* ∈ (0, *T*], we get a system of equations for the solute concentrations *c*_*j*_ for each *j* ∈ *J* = {*e*} ∪𝒫. The system reads

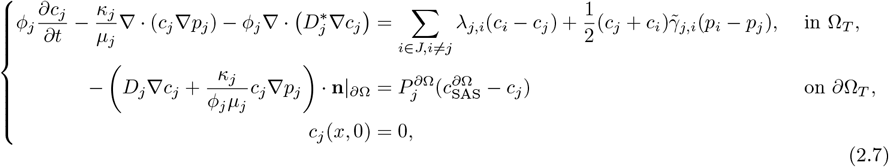

where *κ*_*j*_ is the permeability, *ϕ*_*j*_ is the porosity, *µ*_*j*_ is the fluid viscosity, and 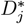 is the effective diffusion coefficient for the compartment *j*. For a membrane separating compartments, we define the diffusive solute transfer coefficient *λ*_*i,j*_ the convective solute transfer coefficient 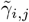,and the osmotic reflection coefficient *σ*_*i,j*_. We also have that 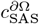 is the concentration in SAS (assumed to be constant in space but dynamic in time and given by the Equation (2.12) defined latter in this paper) and 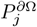 is the diffusive permeability of the boundary layer along the outer surface.

The diffusive and convective solute transfer coefficients are modeled as

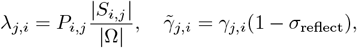

where *P*_*i,j*_ is the permeability of the membrane separating the *i*− th and *j*− th compartments to the solute and *σ*_reflect_ is the solvent-drag reflection coefficient.

### 2.4 Summary of the model

In the rest of this work, we use the multicompartment model with *J* = {*e*} ⋃ 𝒫and 𝒫= {*pa, pv*},ℬ. In (*x, t*) ∈ Ω × (0, *T*), the system reads

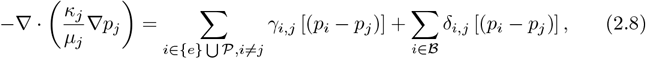

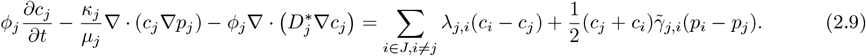

For boundary conditions at the pial surface and at the ventricules surface, *i*.*e*. for (*x, t*) ∈ *∂*Ω × (0, *T*), we have

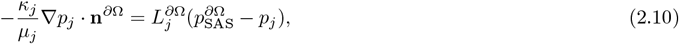

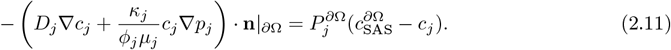

We model intrathecal injection of Gadobutrol. To do so, we assume that the concentration within the SAS is constant and evolves in time following

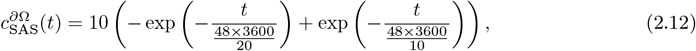

Lastly, we assume no Gadobutrol concentration at the initial state, *i*.*e. c*_*j*_(*x*, 0) = 0, ∀*x* ∈ Ω and for *j* ∈ { *e, pa, pv*}.

Representing the evolution of the concentration of Gadobutrol in SAS for 14 days is given in Fig 1.

**Figure 1:**
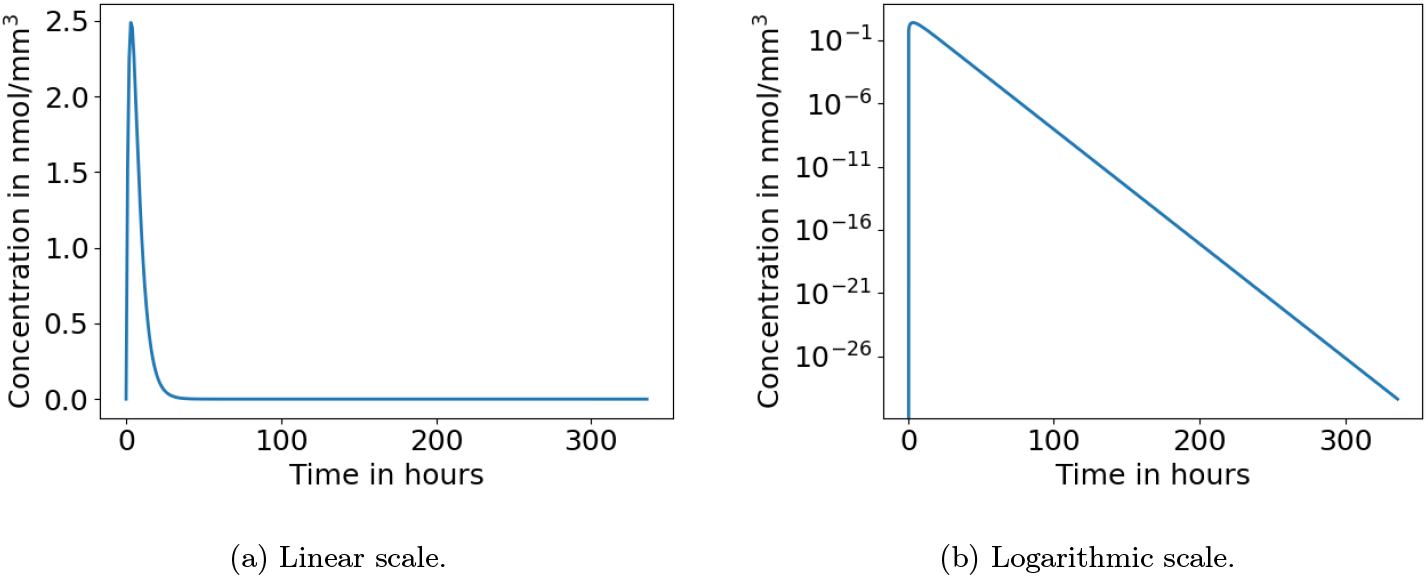
Temporal evolution of SAS tracer concentration after intrathecal injection. This figure is obtained from Equation (2.12).

### 2.5 Computational domain: brain mesh with different zones

To model a brain affected by a tumor, we construct a brain comprising five zones:

1. the gray matter;
2. the white matter;
3. vasogenic peri-tumoral edema;
4. non-vasogenic peri-tumoral edema
5. the tumor.

Hence, we divide our brain domain Ω as

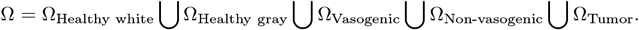

Using “Ernie” brain and meshing techniques from the book [42, 41], we arrive to the brain mesh depicted in Fig. 2. In the following we will consider model variations as well as changes in the mechanical properties in each of the previously presented brain zones to test hypotheses about the disruption of the glymphatic system.

**Figure 2:**
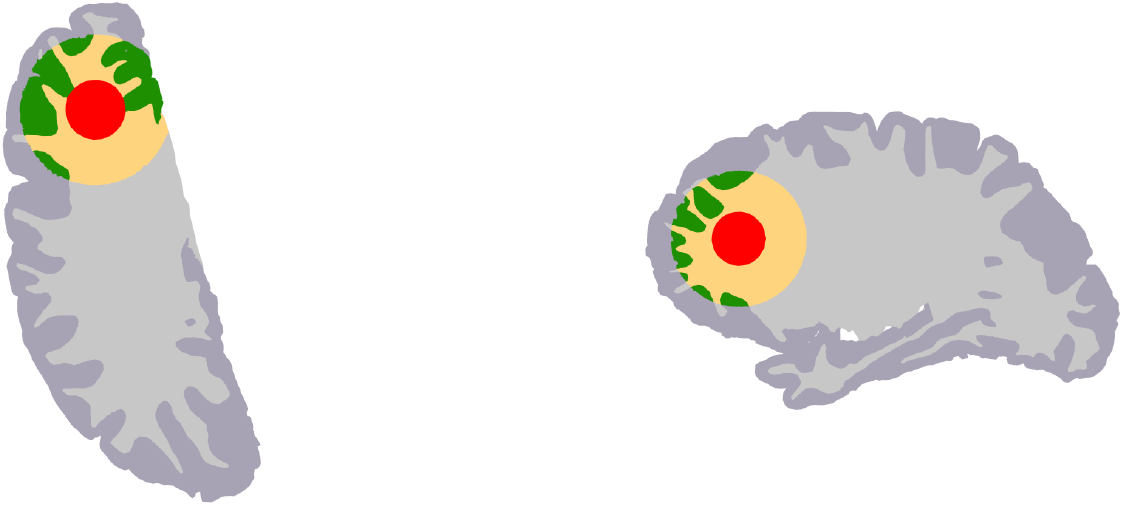
Brain mesh with the 5 zones corresponding to gray matter (gray), white matter (lighter grey), non-vasogenic edema (green), vasogenic edema (yellow) and tumor (red).

As different edema types can develop simultaneously [43] (*e*.*g*. non-vasogenic and vasogenic edema can happen simultaneously and their progressions are linked), we will present in the following a test case that refines the peri-tumoral edema region to consider a zone for the vasogenic part and the non-vasogenic part of the edema.

### 2.6 Parameters values

We present the parameter values in the different compartments of the model.

Table 1 details the symbols, units and definitions for the parameters used in the model. Table 2 summarizes the parameters values and the rest of this subsection describes how to obtain them.

**Table 1:** Definition and unit for each parameter. In this table, *j* ∈ *J* = {*e, pa, pv*}.

| Symbol | Name | Unit |
| --- | --- | --- |
| $\phi_j$ | Porosity | None |
| $D_{\text{free}}, D^*$ | Free and apparent diffusion coefficient | $\text{mm}^2/\text{s}$ |
| $\mu_j$ | CSF dynamic viscosity | $\text{Pa s}$ |
| $\kappa_j$ | Permeability | $\text{mm}^2$ |
| $\lambda_j$ | Diffusive permeability of astrocyte endfeet barrier for gadobutrol | $\text{s}^{-1}$ |
| $\gamma_{e,pa}, \gamma_{e,pv}$ | Hydraulic permeability of astrocyte endfeet barrier | $1/(\text{Pa s})$ |
| $\gamma_{v,pv}$ | Hydraulic permeability of BBB | $1/(\text{Pa s})$ |
| $L_e^{\partial\Omega}$ | Hydraulic conductivity at the pial surface | $\text{mm}/(\text{Pa s})$ |
| $p_j\vert _{\partial\Omega}$ | CSF Pressure at the surface of the brain | $\text{mmHg}$ |
| $P_j^{\partial\Omega}$ | Diffusive permeability at the pial surface | $\text{mm/s}$ |
| $L_j^{\partial\Omega}$ | Hydraulic permeability at the pial surface | $\text{mm}/(\text{Pa s})$ |

**Table 2:** Baseline parameter values. The references indicate the works in which the parameter value is directly found or the works used to compute them. If a * is present next to a reference, it means it was computed using the reference or adjusted using assumptions (see rest of Subsection 2.6 for details). In this table, *j* ∈ *J* = {*e, pa, pv*}.

| Symbol | Value | Reference/computation |
| --- | --- | --- |
| $\phi_j$ | $\phi_e = 0.2, \phi_{pa} = 0.0056, \phi_{pv} = 0.0094$ | [63, 4],* [40, 50] |
| $D_{\text{free}}, D^*$ | $D_{\text{free}} = 3.80 \times 10^{-4}, D_e^* = 1.32 \times 10^{-4},$<br>$D_{pa}^* = 2.13 \times 10^{-6}, D_{pv}^* = 3.56 \times 10^{-6}$ | [62],* |
| $\mu_j$ | $0.7 \times 10^{-3}$ | [7] |
| $\kappa_j$ | $\kappa_e = 2 \times 10^{-11}, \kappa_{pa} = 6.79 \times 10^{-9}, \kappa_{pv} = 6.51 \times 10^{-9}$ | [28],* [49, 20] |
| $\lambda_j$ | $\lambda_{e,pa} = \lambda_{e,pv} = 6.21 \times 10^{-3}$ | * [49, 58] |
| $\gamma_{e,pa}, \gamma_{e,pv}$ | $\gamma_{e,pa} = 1.69 \times 10^{-7}, \gamma_{e,pv} = 1.5 \times 10^{-7}$ | * [49, 64] |
| $\gamma_{v,pv}$ | $7.844 \times 10^{-11}$ | * [49] |
| $L_e^{\partial\Omega}$ | $1.0 \times 10^{-8}$ | * [35] |
| $p_j\vert _{\partial\Omega}$ | $p_{pa}\vert _{\partial\Omega} = 10.03, p_{pv}\vert _{\partial\Omega} = p_e\vert _{\partial\Omega} = 10.0$ | |
| $P_j^{\partial\Omega}$ | $P_{pa}^{\partial\Omega} = P_{pv}^{\partial\Omega} = 3.5 \times 10^{-4}$ and $P_e^{\partial\Omega} = 3.5 \times 10^{-5}$ | * [35] |
| $L_j^{\partial\Omega}$ | $L_{pa}^{\partial\Omega} = 1.2 \times 10^{-7}, L_{pv}^{\partial\Omega} = 1.5 \times 10^{-7}$ and $L_e^{\partial\Omega} = 1.0 \times 10^{-8}$ | * [35] |

#### Porosities

The porosity of the ECS is 0.2 [63]. The volume fraction of the PVS in human white matter is estimated to be around 1.5% [4, 40]. We know the volume distributions for arteries, veins, and capillaries from [50], *i*.*e*.

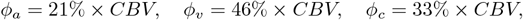

with CBV (for cerebral blood volume) defined as the amount of blood in 100 mL of brain tissue. Relative CBV has no unit as it is a ratio in mL of blood*/*100 mL of tissue.

It’s unclear whether PVSs exist around small capillaries. Furthermore, even if PVS around capillaries exist, it is reasonable to assume that resistance to flow along these structures will be high. In the present work, we assume that the fluid escapes the PVS before reaching the capillaries. Redistributing the porosity used for the PVS around the capillaries in [49], we arrive at the following porosities

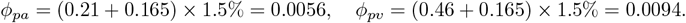

In the previous equations, the factor 0.165 comes from the redistribution of the volume fraction of the capillaries, *i*.*e*. 0.33*/*2 = 0.165.

#### Diffusion coefficients: free and effective

Gadobutrol is a contrast agent molecule commonly used for brain MRI. The free diffusion coefficient of Gd-DTPA (a molecule of 550 Da) is reported in [62] using measurements from [25]. We use the value

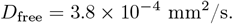

Referring again to [62], we use the tortuosity of the ECS to be *λ*_*e*_ = 1.7. We compute the effective diffusion coefficient using

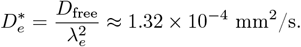

As the tortuosity of the PVS is probably close to 1, effective diffusion the PVS is assumed to be close free diffusion times the porosity of the compartment. Hence, we have

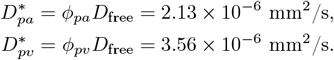

We decided to ignore dispersion effects. It is important to mention that this latter effect is known to contribute to transport of solutes within large PVS. As we represent in this work penetrating vessels, their PVSs are relatively small and we assume no dispersion. The interested reader can find many information about dispersion in PVSs in [2, 21, 51].

#### Tissue permeability and CSF viscosity

We use the permeabilities *κ*_*j*_ (*j* ∈{*e*}⋃ 𝒫) specified in [49]. In this work the authors used the resistance coefficients specified in [64], assumed Poisseuille flow in the PVSs to derive a relation linking the permeabilities *κ*_*j*_ and the resistance coefficients, and used as a reference value the permeability in the ECS given in [28], that is *κ*_*e*_ = 2 × 10^−11^ mm^2^. This led to the values

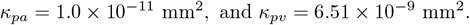

Similar order of magnitudes have been found in [14]. However, it is important to highlight that assuming fluid exits aPVS before the precapillaries, the resistance coefficient indicated in [20] would lead to a permeability coefficient

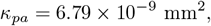

replicating the same computations as in [14] and [49]. Given that it has been observed that the PVS around arteries are larger compared to PVS around veins [56], this latter coefficient appears to be the more reasonable.

Dynamic viscosity of CSF and interstitial fluid is *µ*_*j*_ = 0.7 × 10^−3^ Pa s, (*j* ∈{ *e*} ⋃𝒫) and is given in [7].

##### Remark 1

*It is also worth mentioning recent progresses in quantifying the hydraulic resistance in PVSs. In [9], the authors reported a hydraulic resistance per unit of length* ℛ *to be close to* 2 × 10^6^ *mmHg min/(mL m) (similar order of magnitude is reported for the average of the resistance along the vessel* 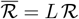 *with L the length of the tube used to perform the approximation of the PVS). This resistance is computed for PVSs around pial arteries, and it is important to stress that resistance at the pial surface is larger due to the elliptic shape of the PVSs. With our units, we have* 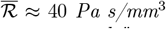, *assuming a characteristic length of L* = 2 *µm (in [9], the authors specified that ℛ is computed “at each normal cross section along the length of the channel by taking the volume-weighted average of the pressure at all elements in 2-µm-thick slices and dividing the pressure difference between adjacent slices by the volume flow rate Q”). Hence, with these new resistance values, the permeability (κ*_*pa*_*) is estimated to be*

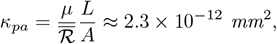

*which is 3 orders of magnitude lower compared to what we use*.

#### Surface area to volume ratio

To compute surface-to-volume ratio of PVSs, we use the surface-to-volume ratio of cerebral blood vessels, estimated to be ≈ 11.77 mm^2^/mm^3^ in [36], and we assume that 1*/*3 × 11.77 mm^2^/mm^3^ corresponds to the surface area to volume ratio for aPVS and we assume the same for surface area to volume ratio for vPVS (1*/*3 is decided arbitrarily to find values close to computations in [19]). We obtain

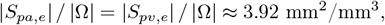

where |*S*_*pa,e*_|represents the total surface area of aPVS and |*S*_*pv,e*_| the one for vPVS, and |Ω| represents the brain volume.

#### Diffusive permeabilities

Diffusive permeabilities between PVSs and ECS are computed using the method specified in [49]. This latter requires to know the Stokes radius of the solute crossing the interface between the compartments. From [58], we know that the hydrodynamic diameter of Gadobutrol (under the commercial name Gadovist) is ≈ 1.0 nm. Thus, we use the Stokes radius *r* = 0.5 nm in our computations to obtain

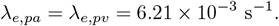

Our coefficients are of the same order of magnitude as the ones found in [35] for penetrating arterioles. Indeed in [35], *C*_*M*_ *D* = 1000 × 3.5 × 10^−7^ mm/s and 3.92 × 3.5 × 10^−4^ ≈ 1.37 × 10^−3^ /s. Hence, our computations give slightly larger diffusive permeabilities compared to [35], but the orders of magnitude are the same.

Using results from [35], we assume that the diffusive permeability at the pial membrane for PVS around arteries and veins is given by

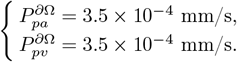

Assuming the pial surface is less permeable to molecules to enter directly the ECS, we assume

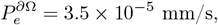

which is one order of magnitude smaller compared to the permeability of aPVS and vPVS at the pial surface.

#### Convective permeabilities

The hydraulic conductivity *L*_*i,j*_ (in mm/(s Pa)) of a porous membrane separating compartments *i* and *j* denotes the ability of the fluid to go through. It depends on the hydraulic resistance of the pores. Previously, in [49], this coefficient was estimated using the resistance coefficients from [64] for the gaps between astrocyte endfeet. The convective permeabilities are defined following

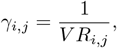

with *R*_*e,pa*_ = 0.57 mmHg*/*mL*/*min and *R*_*e,pv*_ = 0.64 mmHg*/*mL*/*min. Hence, our solute transfer coefficients are

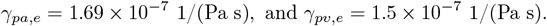

Furthermore, as the contrast agent molecule we consider in this work is small (550 Da for Gadobutrol), we assume that the solvent-drag reflection coefficient is zero, *i*.*e. σ*_reflect_ = 0. This latter assumption was also used in [34]. Therefore, our convective transfer coefficients are defined by

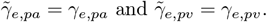

Based on the values reported in [35] for the hydraulic permeability of the astrocyte endfeet barrier, and using the values for the closest layer to the pial surface, we assume

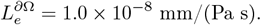

More precisely, we assumed one order of magnitude smaller compared to the values for aPVS and vPVS reported in [35].

We assume for the permeability to the fluid at the pial surface of aPVS and vPVS is infinite, leading to Dirichlet boundary conditions for the pressure fields at the pial surface for these two compartments.

In the following, we will assume in a test case a leakage of fluid from the veins to the vPVS. The blood pressure in veins is assumed to be (estimated in [61])

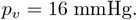

Using the computations described in [49], we estimate (taking two times what we estimated for the murine model)

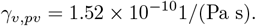

#### CSF pressure in SAS

CSF pressure for humans in the suspinal position is 7 − 15 mmHg [53]. In this work, we take the CSF pressure value to be 10 mmHg. For the pressure at the pial surface for aPVS, we increase this value to apply a pressure difference from aPVS to vPVS. Therefore, we assume a pressure

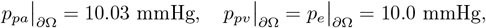

to obtain a pressure difference of similar to the transmantle pressure (estimated to be 4 Pascal in [60]).

### 2.7 Model variations

To study the effect of glioma growth on the glymphatic system, we vary the porosities (*i*.*e*. the volume fractions) of the compartments in the edema and tumor regions. Indeed, permeabilities and diffusion coefficients are linked to the porosities of the compartments.

We know that effective diffusion is related to the tortuosity of the compartment in which the solute diffuses. Indeed, in [62] effective diffusion is given by

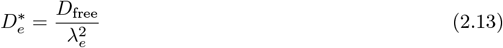

with *λ*_*e*_ being the tortuosity of the ECS. We define a relation that links porosity and tortuosity. We assume a relation of the form

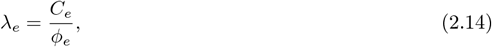

with *C*_*e*_ a constant that will be determined latter.

Looking back to Equation (2.13) given in [62], effective diffusion can be written as

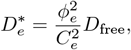

if we assume the extracellular space to be a space filled with spherical inclusions (*i*.*e*. the glial or tumor cells). Using the values for baseline porosity and tortuosity in the ECS, we have

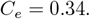

For PVS, the relation that links porosity and tortuosity can not hold as tortuosity is close to one but porosity is very small. Therefore, we assume a simpler relation that links diffusion in the PVS and the porosity, *i*.*e*.

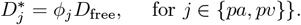

For permeabilities, a well-known equation (see *e*.*g*. [46]) assumes

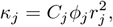

(*j* denoting the compartment) where *ϕ*_*j*_ is the porosity of compartment *j, r*_*j*_ is the radius of the pores and *C*_*j*_ is a constant that is related to the shape of the pores and the tortuosity. Following [5], we assume that this coefficient follows

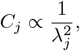

leading to a proportionality relation between permeability and porosity of

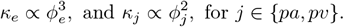

Altogether, in the following, we assume the proportionality relations

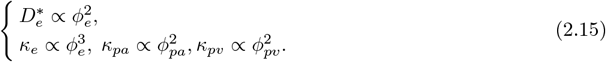

Using these relations, we know that there exist constants 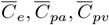 (these constant values must have unit mm^2^ to define permeability correctly) such that

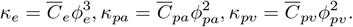

Using this assumption and the values for the baseline porosities and permeabilities, we obtain

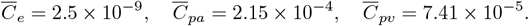

Finally, to study the effect of varying the porosities in the edema or tumor regions, we define a piecewise constant porosity coefficient, *i*.*e*. we assume for each *j* ∈ *J*

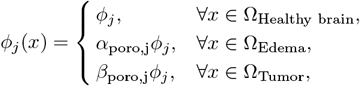

with *α*_poro,j_, *β*_poro,j_. Similarly, we also assume local changes of transfer coefficients between the compartments, *i*.*e. γ*_*i,j*_ and *λ*_*i,j*_ are now defined as piecewise constant functions similarly to the porosities.

#### Remark 2

*Although parameters are piece-wise constant in space, we assume continuity of concentration and pressure at the interface between the sub-regions (healthy tissue, edema and tumor) as well as continuity of the fluxes. This allows to get rid of interface terms in the weak formulation of the model used for finite element simulations*.

In the following section we simulate 1 reference test case with baseline parameter values and 3 test cases corresponding to:

1. **Pure vasogenic edema:** In this test case, we assume that the BBB (of capillaries and veins) is disrupted in the edema region (although it is worth mentioning that in the tumor bed, blood vessels are chaotic and non specific). This leads to a leakage of plasma from the vein compartment to the vPVS. We represent this effect using Equation 2.6. We assume the blood pressure in the veins to be constant and equal to *p*_*v*_ = 16 mmHg. Furthermore, we assume the transfer between vPVS and the veins to be given by

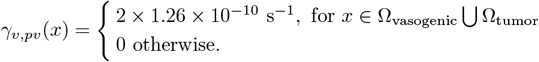 This transfer corresponds to 2 times the transfer coefficient found in [49] between veins and vPVS. We also assume a disruption of the AEF barrier in the edema and tumor region, *i*.*e*. we assume that the fluid transfer coefficients that connect PVSs and ECS are multiplied by a factor 2. Similarly, we assume that the diffusive permeability coefficient of the AEF barrier is multiplied by two. Altogether, for the AEF barrier parameters we have

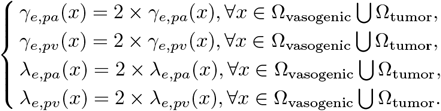 Furthermore, as the pressure from the edema pushes in the ECS, probably enlarging the porosity of the ECS and closing PVSs in the edema region, we assume that the baseline porosity of the ECS in the edema is multiplied by 1.5, *i.e*.

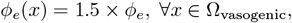

with the *ϕ*_*e*_ on the right-hand side being the baseline porosity given in Section 2.
2. **Pure non-vasogenic edema:** In this test case, we assume that there is no disruption of the integrity of the blood-brain barrier. However, we assume that there is a swelling of the glial cells leading to a reduction of the ECS porosity in the non-vasogenic edema region. Hence, we change the parameter values within the non-vasogenic edema and tumor regions only. To represent the effect of the non-vasogenic edema, we reduce the porosity of the 3 compartments. We therefore assume

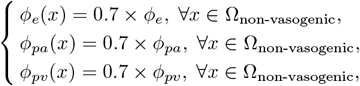

with the *ϕ*_*e*_, *ϕ*_*pa*_, *ϕ*_*pv*_ on the right-hand side being the baseline porosities given in Section 2. To represent the effect of migratory tumor cells, we assume a disruption of the permeability of the astrocyte endfeet barrier separating ECS and vPVS. Indeed, we implicitly assume here that migratory tumor cells tend to follow the direction of the flow (from aPVS to ECS and then vPVS). Thus, in this process, we assume they obstruct the gaps between astrocyte endfeet in the edema (non-vasogenic) and tumor regions, leading to

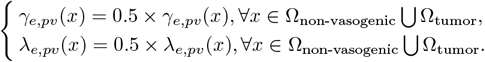
3. **Mixed edema (vasogenic and non-vasogenic):** In this test case, we assume that the edema is of mixed-type. In the gray matter we have a non-vasogenic part and in the white matter we have a vasogenic edema. Hence, we modify parameter values in the two edema regions and in the tumor. The parameter values within the two edemas are chosen following the parameter changes proposed in the two previous test cases: we assume in the vasogenic part of the edema the parameter changes used for the pure vasogenic test case and in the non-vasogenic part of the edema and the tumor, the parameter used in the pure non-vasogenic test case. Furthermore, for the tumor region, we choose baseline porosities and assume that only the fluid and solute transfer coefficients between ECS and vPVS are reduced, leading to

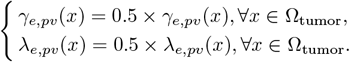

The precise changes of the parameter values compared to the baseline values are indicated in the results section before each test case. We also indicate in Tables 6 and 7 the parameter values used for each test case.

### 2.8 Quantities of interest

In addition to the pressure fields and the spatio-temporal evolution of the concentration of tracer molecules, we compute a quantity to measure the EPR effect. We first define

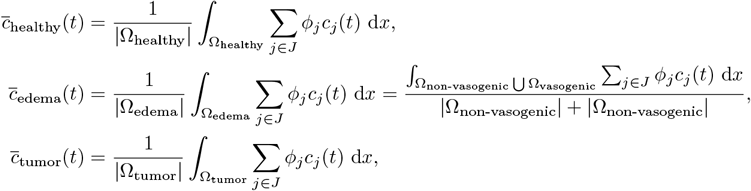

the average tracer concentration in, respectively, the healthy part of the tissue, the edema and the tumor. We also define the total amount (or mass) of solute within the brain

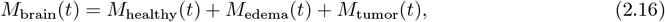

with

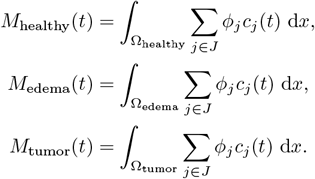

To measure the accumulation of solutes within the ECS in the peri-tumoral and tumor regions, we compute the mean intrinsic concentrations, *i*.*e*.

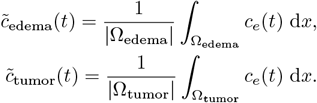

To estimate possible solute retention, we compute ratios (that we name “EPR”) with the reference solution, *i*.*e*.

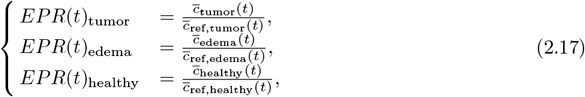

with 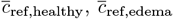 and 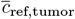 denoting the average concentration of solute in the edema and tumor regions for the reference solution.

To study CSF flow between the compartments and its velocity, we compute the mean velocity of the fluid in the 3 compartments within the brain. This is given by

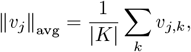

with *K* being the set of mesh nodes. We also investigate the maximum velocity in the brain, given by

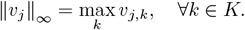

We also compute the debit of CSF transfered between the ECS and aPVS as well as vPVS. This quantity is given using

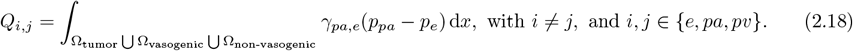

### 2.9 Numerical set-up

We discretize System (2.8)–(2.9) using the finite element method. To implement the scheme, we use the open-source finite element library Fenics [39, 1]. We use P1 finite elements for the pressure and concentration fields. The linear system associated to the finite element discretization of the pressure equations is solved using Conjugate Gradient with “hypre_amg” preconditionning. The linear system associated with the advection-diffusion part is solved using the GMRES algorithms with the ILU preconditioner. Our code is freely accessible at https://github.com/alexandrepoulain/glymphatics-glioma.

## 3 Results

### 3.1 Pressure distributions comparison, CSF flows between the compartments and velocities

Figure 3 presents the pressure fields obtained for the 3 test cases and the reference simulation. Several changes are obtained due to the effects of the edemas. We highlight them in the caption of Figure 3. It is worth mentionning that, as the fluid moves in the opposite direction of the fluid pressure gradient, this disruption of the pressure fields changes locally the direction of the fluid flow within the 3 compartments. However, reduction of the porosities in the tumor region also slows the flow in the tumor region.

**Figure 3:**
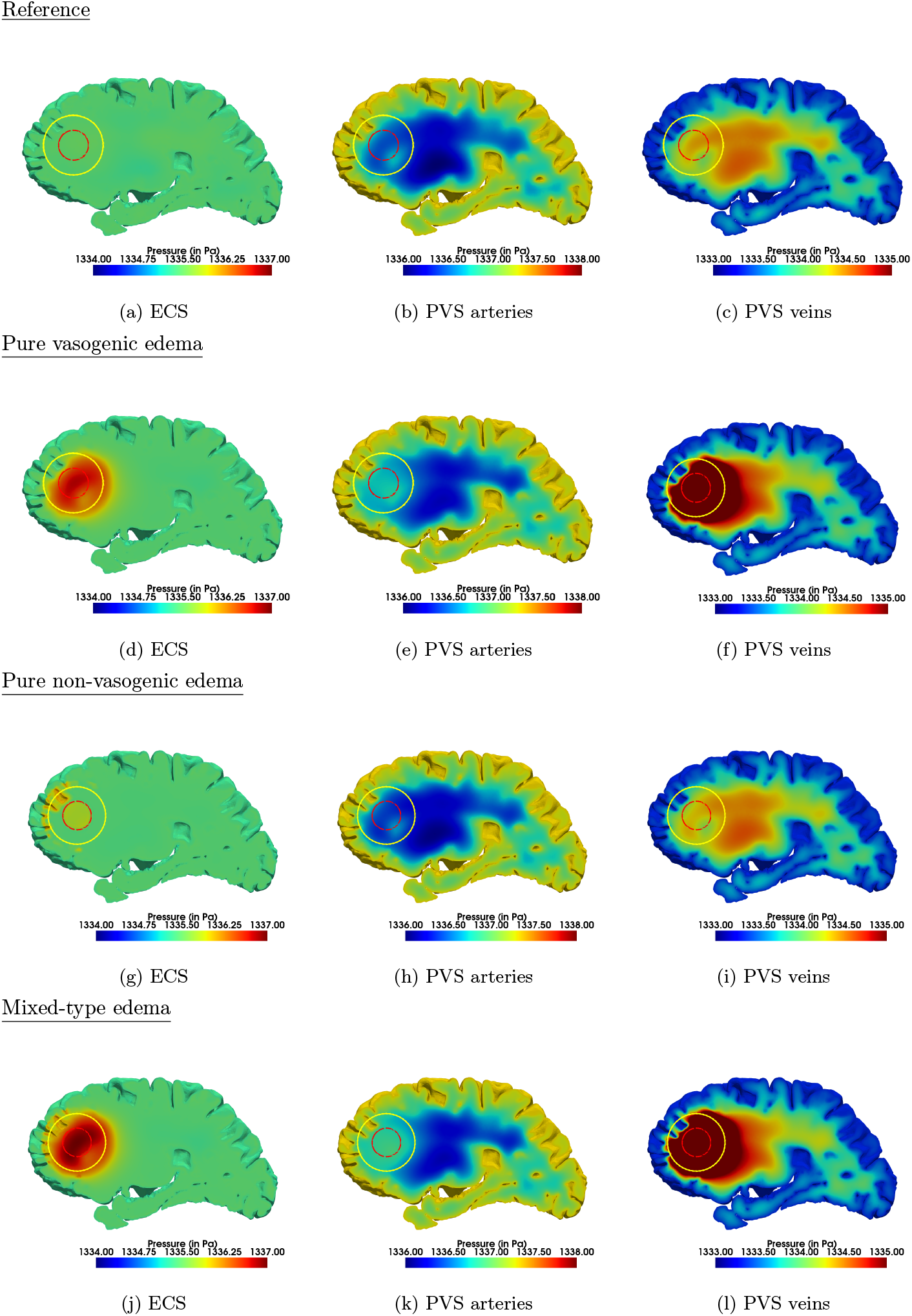
Comparison of pressure fields obtained for baseline parameter values (*i*.*e*. reference) and the 3 test cases. For each image, the yellow circle indicates the boundary of the edema region, while the red circle indicates the boundary of the tumor region. Vasogenic and non-vasogenic edema increase the pressure in the ECS locally in the the tumor and peri-tumor regions compared to reference conditions. A slight pressure disruption in the same regions is also seen in aPVS for the vasogenic edema test case. Furthermore, due to the leakage of fluid from the blood into the vPVS, there is a large increase of pressure in vPVS observed for the pure vasogenic test case. For the mixed-type edema test case, we observe a combination of the effects of both edema types, resulting in a larger pressure increase in the ECS within the tumor and peri-tumor regions compared to the pure vasogenic test case.

To compare the fluid flow between the two test cases presented above and the reference test case, we compute fluid transfer between compartments and the mean as well as the max velocity of the flow. Table 3 gives the debit of the transfer of fluid between the 3 compartments in the edema and tumor regions for the 3 test cases.

**Table 3:**
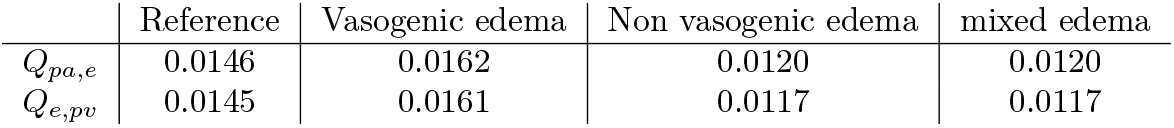
Fluid transfer debit between compartments in (mL/s) within the tumor and edema regions.

#### Remark 3

*We compute the debit using Equation* (2.18) *for the three test cases and the reference even though the region of the edema could vary. Indeed, considering for example the non-vasogenic edema test case, the region of the edema with modified parameter values correspond only to* Ω_*non-vasogenic*_. *However, in order to compare the debit for similar region sizes, we compute the integral over the tumor and the both edema regions regardless of the test case*.

Table 3 reveals that, compared to reference simulation, for vasogenic edema, the fluid transfer between compartments locally in the tumor and edema is enhanced whereas for non-vasogenic edema, the fluid exchange is decreased. For mixed-type edema, the fluid exchange is slightly decreased compared to the reference case, an corresponds to the same transfer as for the non-vasogenic edema.

Table 4 summarizes the mean and maximum velocity of the fluid for the three compartments in domain (*i*.*e*. the brain) and for the three test cases.

**Table 4:** Mean and maximum fluid velocity in the brain for the three test cases (in mm/s).

|  | Reference | Vasogenic | Non-vasogenic | Mixed-type |
| --- | --- | --- | --- | --- |
| $\ v_e\ _{\text{avg}}$ | $1.43 \times 10^{-8}$ | $1.70 \times 10^{-8}$ | $1.65 \times 10^{-8}$ | $1.88 \times 10^{-8}$ |
| $\ v_{pa}\ _{\text{avg}}$ | $1.40 \times 10^{-4}$ | $1.42 \times 10^{-4}$ | $1.39 \times 10^{-4}$ | $1.37 \times 10^{-4}$ |
| $\ v_{pv}\ _{\text{avg}}$ | $8.49 \times 10^{-5}$ | $1.14 \times 10^{-4}$ | $8.31 \times 10^{-5}$ | $1.14 \times 10^{-4}$ |
| $\ v_e\ _{\infty}$ | $5.87 \times 10^{-7}$ | $5.70 \times 10^{-7}$ | $5.73 \times 10^{-7}$ | $5.69 \times 10^{-7}$ |
| $\ v_{pa}\ _{\infty}$ | $1.58 \times 10^{-3}$ | $1.43 \times 10^{-3}$ | $1.52 \times 10^{-3}$ | $1.43 \times 10^{-3}$ |
| $\ v_{pv}\ _{\infty}$ | $8.05 \times 10^{-4}$ | $2.42 \times 10^{-3}$ | $9.24 \times 10^{-4}$ | $3.34 \times 10^{-3}$ |

**Table 5:** List of frequently used acronyms.

| Acronym | Meaning |
| --- | --- |
| CNS | Central nervous system |
| CSF | Cerebro-spinal fluid |
| ADC | Apparent diffusion coefficient |
| EPR | Enhanced permeability and retention |
| BBB | Blood-brain barrier |
| CBB | CSF-blood barrier |
| ECS | Extra-cellular space |
| SAS | Subarachnoid space |
| PVS | Peri-vascular space |
| vPVS | Peri-vascular space around the veins and venules |
| aPVS | Peri-vascular space around the arteries |

**Table 6:** Parameter values used in the vasogenic edema region. We use baseline parameter values in the vasogenic edema region for the parameters not indicated in this Table. In this table, *j* ∈ *J* = {*e, pa, pv*}.

| Symbol | Value |
| --- | --- |
| $\phi_j$ | $\phi_e = 0.3, \phi_{pa} = 0.0056, \phi_{pv} = 0.0094$ |
| $D_j^*$ | $D_e^* = 2.96 \times 10^{-4}, D_{pa}^* = 2.13 \times 10^{-6}, D_{pv}^* = 3.56 \times 10^{-6}$ |
| $\mu_j$ | $0.7 \times 10^{-3}$ |
| $\kappa_j$ | $\kappa_e = 6.75 \times 10^{-11}, \kappa_{pa} = 6.79 \times 10^{-9}, \kappa_{pv} = 6.51 \times 10^{-9}$ |
| $\lambda_j$ | $\lambda_{e,pa} = \lambda_{e,pv} = 1.24 \times 10^{-2}$ |
| $\gamma_{e,pa}, \gamma_{e,pv}$ | $\gamma_{e,pa} = 3.37 \times 10^{-7}, \gamma_{e,pv} = 3.01 \times 10^{-7}$ |
| $\gamma_{v,pv}$ | $7.844 \times 10^{-11}$. |

**Table 7:** Parameter values used in the nonvasogenic edema region. We use baseline parameter values in the vasogenic edema region for the parameters not indicated in this Table. In this table, *j* ∈ *J* = {*e, pa, pv*}.

| Symbol | Value |
| --- | --- |
| $\phi_j$ | $\phi_e = 0.14, \phi_{pa} = 0.0039, \phi_{pv} = 0.0066$ |
| $D_j^*$ | $D_e^* = 6.44 \times 10^{-5}, D_{pa}^* = 1.50 \times 10^{-6}, D_{pv}^* = 2.49 \times 10^{-6}$ |
| $\kappa_j$ | $\kappa_e = 6.86 \times 10^{-12}, \kappa_{pa} = 3.33 \times 10^{-9}, \kappa_{pv} = 3.19 \times 10^{-9}$ |
| $\lambda_j$ | $\lambda_{e,pv} = 3.11 \times 10^{-3}$ |
| $\gamma_{e,pv}$ | $\gamma_{e,pv} = 7.51 \times 10^{-9}$ |

Table 4 indicates that, for our three test cases, the average velocity in the ECS slightly increases compared to the reference case, with the most important increase for the mixed-type edema. Interestingly, the opposite is observed for the maximum velocity in the ECS. For the aPVS, the average and maximum velocity do not seem to be affected by the edema in the three test cases compared to the reference case. However, fluid velocity in the PVS around veins are modulated due to the effects of the edemas. For the vasogenic edema test case, an increase in fluid velocity (maximum and mean) in the PVS veins compartment compared to reference is observed for the vasogenic and mixed-type edema test cases. This effect is probably due to the perfusion of fluid from the blood compartment to the PVS veins, hence disrupting the pressure field in this compartment and leading to an increase in velocity.

### 3.2 Comparison of tracer concentration evolution and clearance

We compare the spatio-temporal evolution of the concentration of Gadobutrol at 4 different time points after simulated intrathecal injection: *t* = {12*h*, 24*h*, 48*h*, 168*h*. After *t* = 168*h*}, the remaining solute mass is slowly cleared and no additional observation could be made compared to the snapshot at *t* = 168*h*.

Figure 4 depicts these 4 snapshots for the 3 test cases and the reference simulation. Comparing the concentration fields for the different cases, we clearly observe that the changes of the porosity parameter creates a disruption of the uptake and clearance of Gadobutrol. Two main observations can be made. First, the vasogenic edema has the effect to fasten the transport of Gadobutrol in the edema region. This is due to the increased porosity resulting in a larger diffusion in this region. As a result, a retention of Gadobutrol in the peri-tumor region is observed even 7 days intrathecal injection. Second, the non-vasogenic edema reduces slightly the transport of Gadobutrol within the brain. Indeed, as seen at *t* = 24*h* for the pure vasogenic test case, the non-vasogenic edema appears as a region with a very small concentration (the non-vasogenic edema corresponds to the small darker regions in the gray matter). This is due to the reduced porosity in this region for this test case, resulting in a smaller macroscopic concentration, computed for example for the ECS compartment with *ϕ*_*e*_(*x*)*c*_*e*_(*x*). For the mixed type edema test case, we mostly see on Figure 4 the effect of the vasogenic edema part. However, although convenient to represent spatially the evolution of Gadobutrol during the simulations, Figure 4 does not allow to compare locally the amount of tracer in specific regions of the brain for the 3 test cases and the reference.

**Figure 4:**
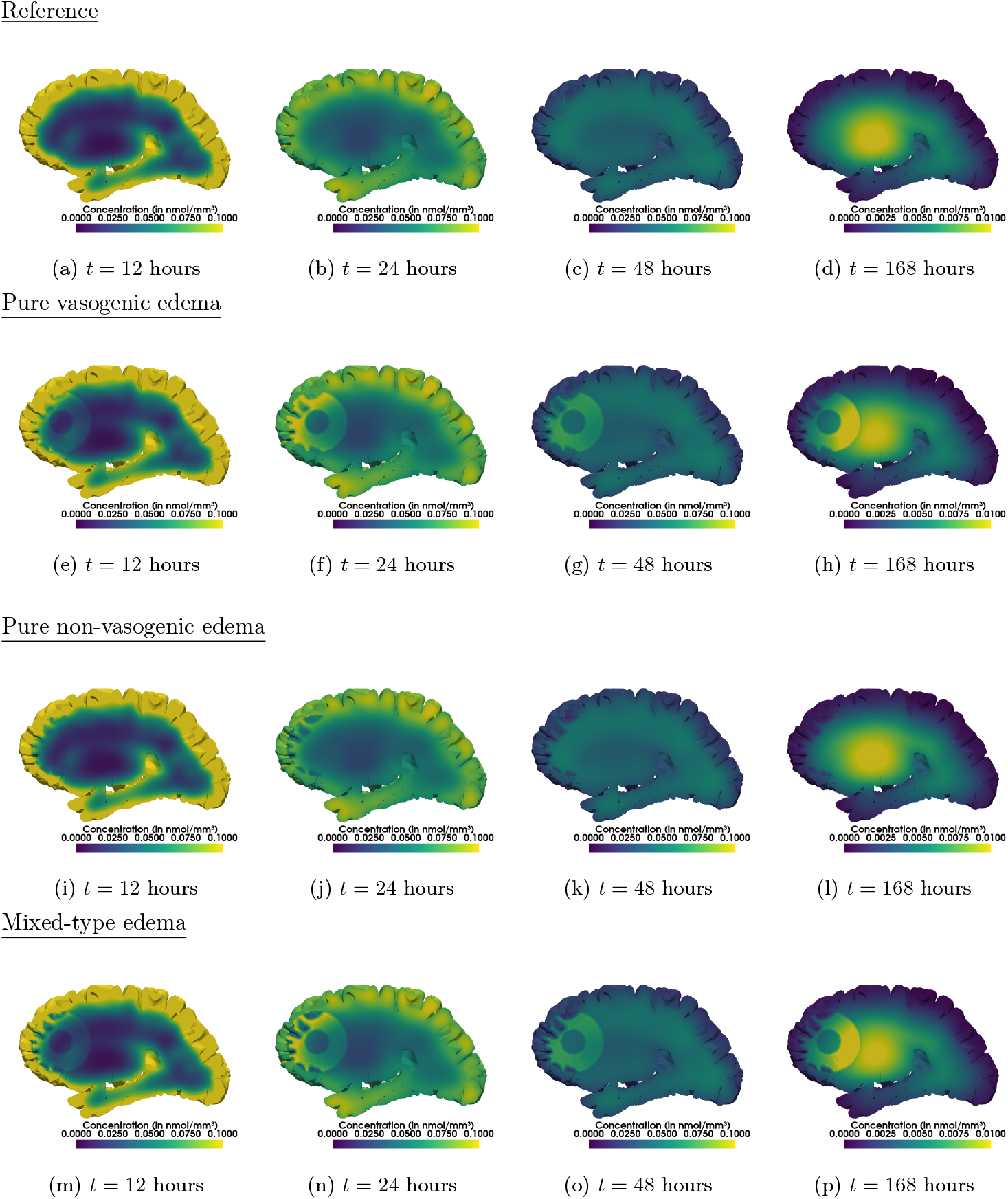
Spatio-temporal evolution comparison of the concentration of Gadobutrol in the brain. Furthermore, the vasogenic edema results in a retention of Gadobutrol in the peri-tumor region observed even at *t* = 168 hours. We observe that the non-vasogenic edema localized in the gray matter (hence close to the surface of the brain) slows the transport of the solute in the depth of the brain. This is seen at 24 hours after injection, where we see that the zones with the smaller concentration of Gadobutrol in the gray matter correspond to the non-vasogenic edema. In this case, the edema would be clearly identified on T1-weighted MR images as it will appear as a darker region. For the mixed-type edema, after simulated injection of Gadobutrol, we observe a build-up of tracer within the edema region. Careful inspection of the concentration distribution within the edema reveals that the increase is mostly localized within the vasogenic part of the edema.

To evaluate clearance speeds within the brain for our three test cases and the reference case we start by inspecting the temporal evolution of the mass of Gadobutrol in the healthy part of the brain, in the edema region and the tumor. Figure 5 depicts the temporal evolution of the tracer mass in the total left hemisphere (*i*.*e*. our computational domain). Very small differences could be observed between the three test cases and the reference computing the evolution of the total mass of tracer within the brain. To compare clearance of solutes within the brain and especially in specific regions of interest such as the edema and the tumor, we use the EPR ratios defined in Equation (2.17).

**Figure 5:**
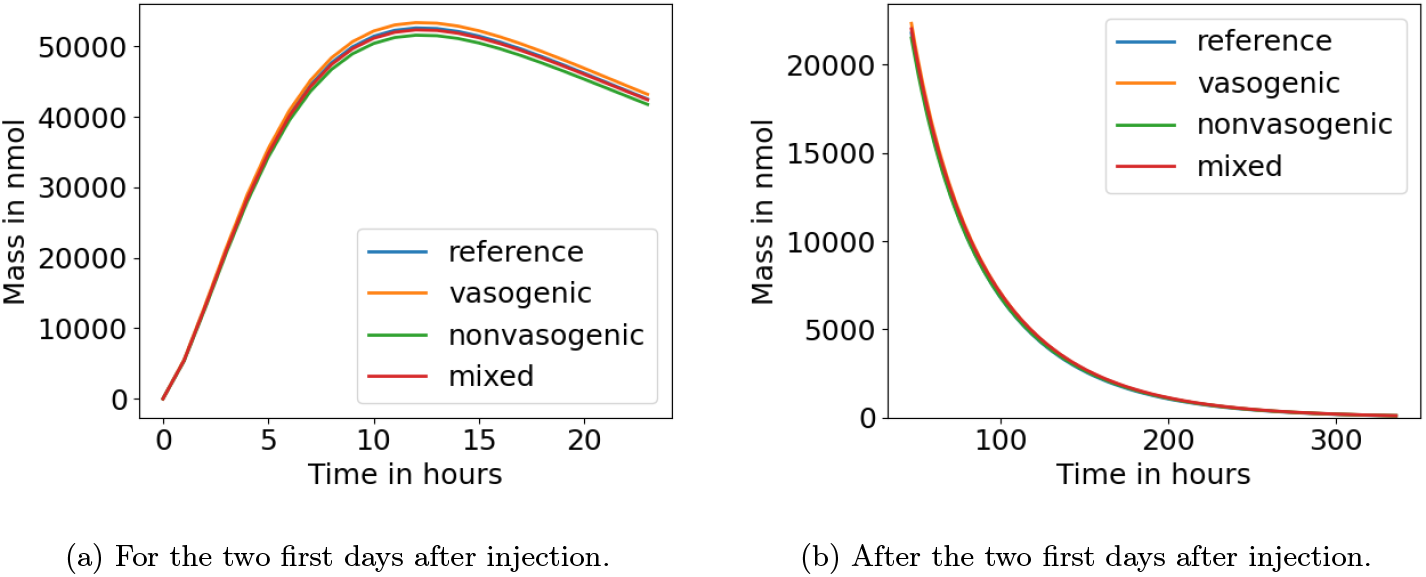
Temporal evolution of total mass of tracer in the healthy part of the brain for the four simulations.

Figure 6 depicts the temporal evolution of the EPR ratio within the tumor and the edema regions.

**Figure 6:**
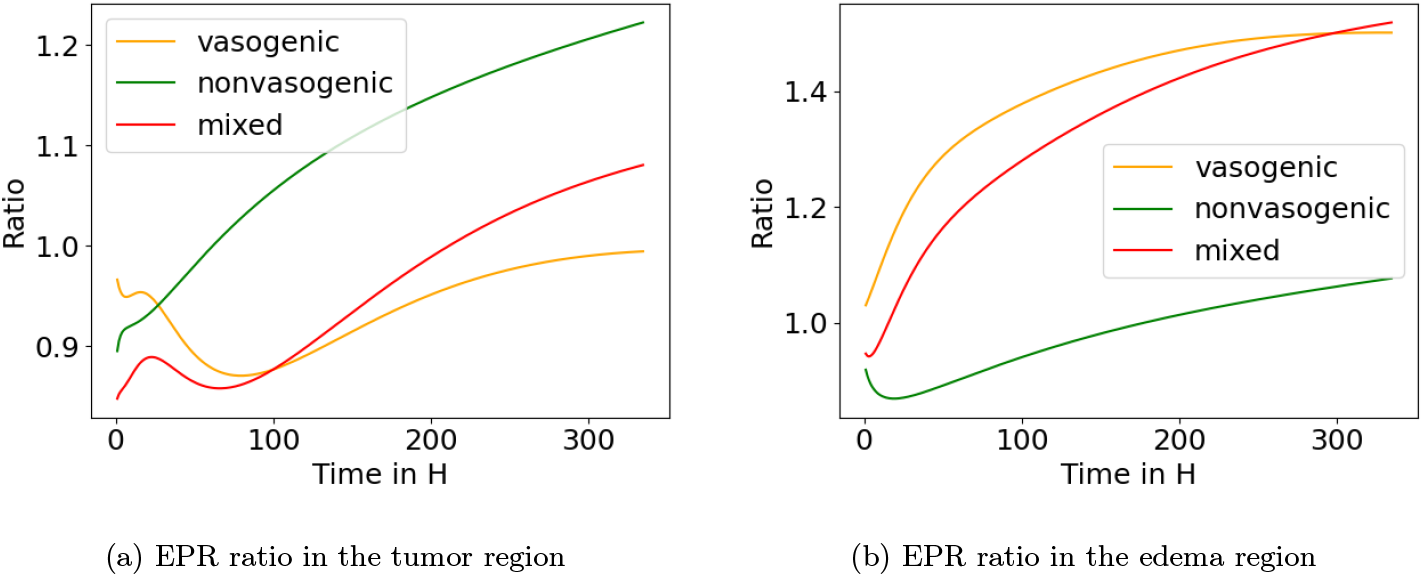
Temporal evolution of the EPR ratio computed using Equation (2.17) for the edema and tumor regions. We observe that at the beginning of the simulations, the mass in the tumor region is smaller compared to the reference for the three test cases. Around the 6th day after injection, the disruption of the glymphatic system produces a retention effect and the mass within the tumor region becomes larger compared to reference case. Around 200 hours after injection, the mass within the tumor for the non-vasogenic test case is above 10% larger compared to reference and for the mixed-type edema the retention effect begins. At the end of the simulation the mass within the tumor for the non-vasogenic test case is 20% larger compared to the reference case. For the vasogenic test case, clearance in the tumor region appears faster. Within the edema region, the mass of tracer is larger compared to the reference from the beginning of the simulations for the vasogenic and mixed-type edema. This is due to the inflated ECS for these two test cases (*i*.*e*. ECS porosity is assumed to be larger in the vasogenic edema). The amount of tracer increases compared to the reference for this two test cases throughout the simulations and approximately 45% more tracer are obtained in the edema region compared to the reference for the vasogenic and mixed-type edema test cases.

Figure 7 shows the temporal evolution of the EPR ratio within the entire domain (*i*.*e*. the left brain hemisphere).

**Figure 7:**
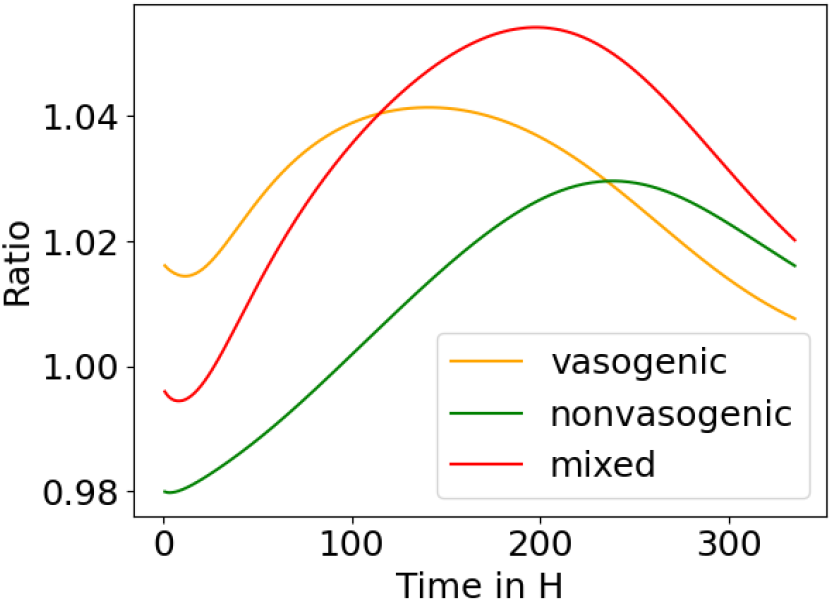
Temporal evolution of the EPR ratio for the total hemisphere. We observe that the three test cases lead to different disruptions of clearance depending on the edema type. For the mixed-type edema, the magnitude of the clearance disruption appears to be the largest. For this test case, there is a approximately 5% more solute in the domain at 200 hours after injection of tracer.

## 4 Discussion

In this work we have seen that peri-tumoral edema causes an impairment of the glymphatic system and a re-distribution of the tracers delivered by intrathecal injection. We demonstrate using simulations that that the type of edema is crucial for whether there is enhanced or impaired delivery. In the vasogenic edema, the delivery is more than 20% higher starting from 8 hours after intrathecal injection of tracer. On the other hand, in the non-vasogenic tracer delivery appears to be vary small. This edema type also appears to be a barrier to the transport of the tracer outside the brain. Indeed, as non-vasogenic edema affects the gray matter, tracer molecules trying to diffuse outside the brain (from the depth to the cortical surface) are slowed due to the non-vasogenic edema, thus leading to retention effect of more than 10% of tracer mass in the tumor region compared to normal conditions. As such, imaging based on intrathecal contrast combined with modeling may give insight into the underlying mechanisms of glymphatic clearance impairment depending on tumor type.

We detail below the observation of the different processes.

### Vasogenic edema enhances tracer delivery in the edema region following intrathecal injection, disrupts the glymphatic system and leads to a retention in the peri-tumor region

Vasogenic edema is a consequence of leaky blood vessels in the tumor and peri-tumor region that are produced due to inflammatory and angiogenic factors released by the tumor [57]. This excess of fluid in the brain tissue creates a local region of large pressure that is well captured by the model (see Figure 3). Under such pressure the brain tissue deforms and probably increases the pore size within the ECS. This latter assumption follows the fact the Apparent Diffusion Coefficient measured by MR images is increased in the peri-tumor region of aggressive tumors indicating the presence of freer stagnant fluid in this region [33]. Assuming these effects and simulating with our model an intrathecal injection of Gadubutrol, we observe an interesting effect: an increase of the number of molecules within the peritumor region (*i*.*e*. the region in which the porosity of the ECS is increased). This observation is due to two effects: the increased ECS volume and, consequently, the enhanced diffusion coefficient in the peri-tumor region region. This leads to a well-defined difference between the edema and the rest of the brain. This difference appears very clear between 2 and 3 days after injection of the tracer as seen on Figure 4. It is also important to mention that the model also predicts a more important transfer of fluid between the PVSs and the ECS in the edema and tumor region compared to a reference solution obtained using baseline parameter values. Furthermore, the mean fluid velocity in the PVS around veins appears to be increased in the context of a vasogenic edema. Altogether, our simulation results indicate that the porosity changes combined with pressure changes due to the leakage of fluid from blood vessels lead to an intensity increase that allows to clearly identify the edema region two days after intrathecal injection of a tracer. Furthermore, these changes due to the edema lead to an enhancement of the glymphatic system with an increased mean fluid velocity in the PVS around veins and an increase of fluid transfer between PVSs and the ECS. This could contribute to the increase of the apparent diffusion coefficient often observed in the vasogenic edema region of a tumor.

### Non-vasogenic edema due to cell swelling produces retention of solute tracer in the tumor region

One of the most important cause of non-vasogenic edema in the brain is the swelling of glial cells [44]. This reduces the ECS volume in the edema region. Inspecting Equations (2.15), we understand that this diminution of the pore size leads to a reduction in the permeability and the diffusion coefficients. Hence, solute and fluid movement are expected to be reduced in this region. Furthermore, assuming that the swelling of the cells combined with the growth of the tumor and possible migration of cells in the PVS around veins, simulations predict an increase of fluid pressure in and around the tumor. This pressure increase is due to a difference of permeabilities and fluid transfer between PVS around arteries and PVS veins. Indeed, as we assume that PVS around veins are impacted by the possible migration of tumor cells, porosity and astrocyte endfeet barrier separating PVS veins and ECS are modulated and produce a trap of fluid in the ECS. Hence, our model indicates that large local pressure close to the tumor and with an intact BBB could be explained by the migration of tumor cells in the PVS combined with the swelling of astrocytes, thus, blocking the fluid to exit the brain. This appears clearly in Table 3. We observe a drop of fluid exchange in the tumor and edema region between PVSs and the ECS. This indicates that non-vasogenic edema reduces fluid convection across compartments. However, it is worth mentioning that there is a slight increase in fluid velocity within the ECS for the non-vasogenic edema compared to the reference simulation (see Table 4).

Simulations also predict that non-vasogenic edema impact the diffusion and clearance of a tracer following intrathecal injection. Indeed, the macroscopic concentration (*i*.*e*. the concentration that is accessible using MR images) decreases in the edema and tumor region (see Figures 4). This observation is explained by the reduced porosity in the ECS due to the swelling of the cells. However, it is interesting to note that our model predicts that the tracer will still reach the tumor as the spatially reduced ECS in the edema region is bypassed by the PVS around arteries. Thus, the tumor is still perfused by the PVSs and will collect tracer molecules. Furthermore, Gadobutrol appears to be trapped in the tumor as it needs to cross the edema in which solute movement is reduced to be cleared out of the brain. Also, the secondary route of clearance which are the PVS around veins are impacted in our scenario, hence, trapping the solute in the tumor region.

Therefore, our results clearly suggest that the resulting impairment from non-vasogenic edema and the migration of tumor cells lead to solute build-up and disruption of the clearance mechanisms locally.

### Biologically relevant test case combining non-vasogenic and vasogenic edemas in the tumor and peri-tumor regions indicates an enhancement of the retention effect

Indeed as indicated by our simulations, simulating a mixed-type edema in which non-vasogenic edema affects the gray matter and vasogenic edema affects the white matter leads to a enhanced retention effect compared to the pure non-vasogenic test case. One explanation for this effect is the following: as vasogenic edema gathers more tracer molecules and is located within the white matter, more solute molecules have to cross the non-vasogenic edema to be cleared our of the brain from its surface. As the non-vasogenic edema acts as a barrier and PVS around veins are blocked by migratory tumor cells, clearance is slowed in the tumor and edema regions. Furthermore, we assumed a reduced fluid and solute transfer between the ECS and vPVS to represent the potential effect of migratory tumor cells blocking the gaps between astrocyte endfeet. This assumption resulted in a reduced fluid transfer between the PVSs and the ECS. This could also partially explain the retention of solutes as less convective movement between the compartments is obtained in our simulations. Hence, our simulations indicate how in certain biological contexts, solutes could be trapped in and around the tumor.

### Perspectives for imaging and therapy

These predicted retention effects of solutes in tumor region following intrathecal injection open the door to possible therapeutic perspectives. Indeed, macroscopic retention of tracer molecules can be estimated from MR images as it corresponds to an increase of intensity on T1-weighted images. An increase of intensity in specific regions that would correspond to an accumulation of tracer molecules in the parenchyma is predicted, by our model. This observation is due to the increase of ECS porosity from fluid leakage originating from blood vessels. This observation from numerical simulations aligns well with observations of the intensity increase on T2 or FLAIR MRI images associated with vasogenic edema. Therefore, the model suggests that the detection of vasogenic edema can be confirmed following intrathecal injection of Gadobutrol with a larger difference between the edema and the normal tissue for images acquired between 2 and 4 days after injection.

Potential therapeutic strategies can also be discussed from our model predictions. Due to property changes made by non-vasogenic edema, a retention of tracer molecules following intrathecal injection is observed due to the local impairment of clearance mechanisms. This effect could be used to inject to the patient a treatment that targets the tumor. Chemically activated or delayed activation chemotherapies [66] could be used to target tumor cells. Indeed, from intrathecal injection, hence bypassing the blood-brain barrier, the retention of the molecules in the tumor region to local glymphatic impairment that is predicted by the model due to the non-vasogenic edema could help to target the tumor cells and reduce the side effects of a systemic treatment.

Therefore, our model highlights two possible glioma-induced glymphatic system alterations that could be used to collect better images and to treat the tumor using a delayed activation chemotherapy taking advantage of the retention of solutes.

### Limitations and further works

Although our results allow to give a possible explanation of the glymphatic system changes due to glioma, our model is currently a simplification of the real problem. Indeed, our modeling assumes that the tissue is static and the boundary of the domain does not deform from the edema pressure. We still implemented porosity changes that could occur within the tissue for the different edema types, but the fact that the brain is not deformable is not consistent with biological observations. Inspired by the work in [59], we are working toward adapting our multicompartment model to include possible brain tissue deformations.

Of primary importance for the potential therapeutic approaches, it is worth mentioning that macrophages in the vicinity of the tumor, which could participate to 30 to 50% of the tumor mass [29], are probably responsible in part to the retention of molecules in this region. However, our work indicates that the impairment of the glymphatic system due to changes in mechanical properties induced by the edema could also contribute to a retention of solutes in the tumor region.

## 5 Conclusion

Our numerical results show the ability of the model to represent the possible glioma-induced impairment of the glymphatic system. Simulating biologically relevant changes of parameter values associated to the effects of the edemas in the peri-tumor region, we identified fluid flow changes in and between the ECS and PVSs. Furthermore, our numerical results also showed an alteration of the clearance of Gadobutrol in the tumor and peri-tumor regions depending on the edema subtype (*i*.*e*. vasogenic, non-vasogenic or combined edema types). Altogether, our results align well with the reduced glymphatic system activity recently observed in patients suffering from gliomas [24]. Based on numerical simulations, we are able to draw potential perspectives for imaging and treatments of glioma.

## 6 Acknowledgments

A.P. acknowledges the support of the CDP C2EMPI, as well as the French State under the France-2030 programme, the University of Lille, the Initiative of Excellence of the University of Lille, the European Metropolis of Lille for their funding and support of the R-CDP-24-004-C2EMPI project. K.-A.M. acknowledges funding and support from Stiftelsen Kristian Gerhard Jebsen via the K. G. Jebsen Centre for Brain Fluid Research, the Research Council of Norway via grants #300305 (SciML) and #301013 (Alzheimer’s physics), from the national infrastructure for computational science in Norway, Sigma2, via grant #NN9279K, and the European Research Council under grant 101141807 (aCleanBrain). K.E.E. acknowledges funding and support European Union’s Horizon 2020 Programme: ERC Grant Agreement No. 758657-ImPRESS, Helse Sør-øst Regional Health Authority grants 2023050, 2021057, 2017073, the Research Council of Norway grants 325971, 261984

## A Formal derivation of the equations

Defining a brain wide clearance model is inherently an upscaling problem. The Glymphatic system explains fluid movement and solute transport at the microscopic scale. However, a brain-wide clearance model can not resolve directly the microscopic effects (this would require to mesh the brain at the scale of the cells and this will be too expensive from a numerical point of view). However, thanks to mathematical techniques of homogenization, we can derive a set of equations representing accurately the glymphatic system at the macroscopic scale.

### A.1 Volume averaging technique

To obtain our brain-wide solute transport and fluid flow model, we first describe the problem at the microscopic scale (the scale of the cells and the PVSs). From there, we seek an average macroscopic description that represents accurately our problem while being computationally less expensive to solve compared to the microscopic problem. To do so, we rely on the volume averaging technique (see *e*.*g*. [68]). It is worth mentioning that this is not the only method to upscale microscopic equations as multiscale asymptotics homogenization produce similar, yet different on several aspects, results [13].

We here define the tools that we will use in the derivation. We define the averaging volume *V* (*x*) Ω ⊂ (Ω being the computational domain, *i*.*e*. an open bounded set of ℝ^3^ with a smooth boundary) as a closed set defined at the point *x*. From a modelling point of view, Ω is the brain. We will refer to the volume of *V* with |*V* | ≡ ∫_*V*_ 1 d*ξ*. Within the averaging volume, we assume that we have multiple compartments *V*_*j*_ (with *j* ∈ *J, J* being the set of indices for the different compartments), such that ⋃ _*j*_ *V*_*j*_ = *V*. In the present work we assume that fluid and solute move within the ECS (*e*), the PVS around arteries including arterioles (*pa*) and the the PVS around veins including venules (*pv*). Other compartments are considered but are represented as impermeable for solute and fluid: *i*.*e*. the blood compartments (veins *v* and arteries *a*) and the glial cells (*c*). We illustrate the volume *V* and its components in Figure 8.

**Figure 8:**
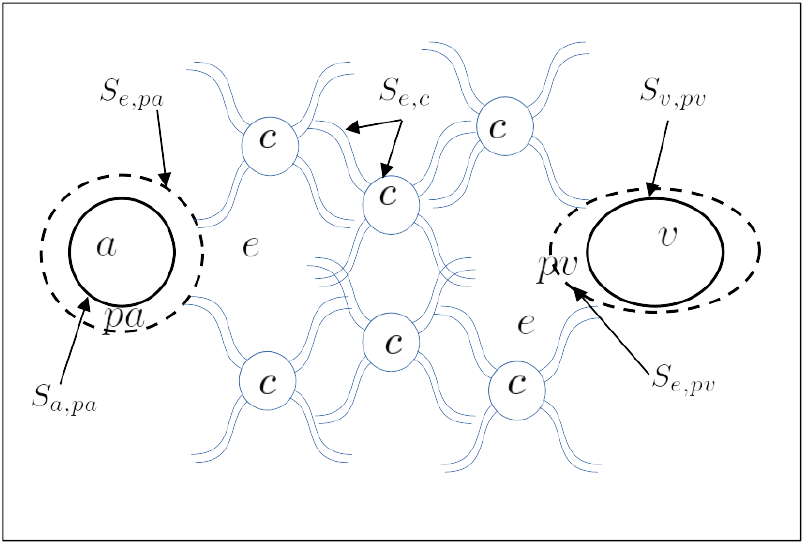
Illustration of the averaging volume. Partial volumes *V*_*j*_ are indicated using their indices (*e* for ECS, *c* for cells, *a* and *pa* for arteries and PVS around them, *v* and *pv* for veins and PVS around them).

The surface of *V*_*j*_ is defined as the surface shared with *∂V* (the outer boundary) and the surface *S*_*j*_ that separates the compartment *V*_*j*_ from other compartments within the volume *V*. In practice, we break this latter surface into the union of the surfaces *S*_*j,i*_ separating compartment *j* and the other compartments *i* ∈ *J* \ *j, i*.*e. S*_*j*_ = ⋃ _*i*_ *S*_*j,i*_. We mention here that from a modeling point of view the surfaces *S*_*j,i*_ are 2-dimensional manifolds that represent the membranes separating the compartments within the brain. These membranes are obviously 3 dimensional objects in nature but since their thickness is very small we collapse them into surfaces. As a consequence, the quantities in our model such as concentration fields are expected to be discontinuous across these surfaces and we will rely to Kedem-Katchalsky equations [31] to describe the fluxes across these.

We define the moving volume average as

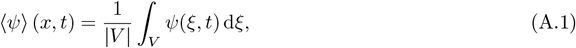

with *ψ*(*ξ, t*) any scalar, vector or second-order tensor quantity. As our goal system is a multicompartment model, we need to define the compartment average (as for multiphase systems)

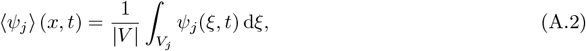

with *V*_*j*_ being the portion of *V* occupied by the compartment *j*. We also define the intrinsic average

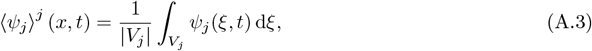

and we have

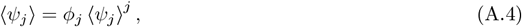

with 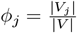 being the porosity (*i*.*e*. the volume fraction) of the compartment *j*.

To upscale our microscopic model, we will use three theorems that can be found in [26].

#### Theorem 4

*For a sufficiently smooth vector field* **T**_*j*_, *we have*

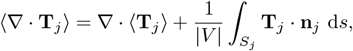

*with* **n**_*j*_ *the unit outward pointing normal to the surface S*_*j*_.

#### Theorem 5

*For a sufficiently smooth scalar field T*_*j*_, *we have*

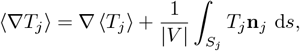

#### Theorem 6

*For a sufficiently smooth scalar field T*_*j*_, *we have*

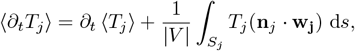

*with* **w**_**j**_ *being the velocity of the interface S*_*j*_.

In the latter theorem, the velocity of the interface *S*_*j*_ appears. As we assume the brain to be static (no displacement of the structures within the brain), the velocity **w**_**j**_ = 0 and the second term on the right-hand side of Theorem 6 cancels out, *i*.*e*. ⟨*∂*_*t*_*T*_*j*_⟩ = *∂*_*t*_ ⟨*T*_*j*_⟩.

### A.2 Microscopic scale model

We first define solute transport and fluid movement at the microscopic scale. We denote by *ρ*_*j*_ the density of the fluid in the compartment *j*, **v**_*j*_ its velocity and *c*_*j*_ the concentration of the solute in compartment *j*. For *j* ∈ {*e, pa, pv*}, the mass balance equation for the fluid reads (for *t* ∈ (0, *T*))

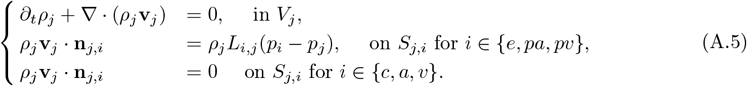

For *j* ∈ {*e, pa, pv*}, the momentum equation reads (for *t* ∈ (0, *T*))

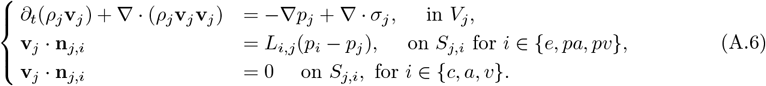

with *L*_*i,j*_ the hydraulic conductivity of the membrane separating compartments *i* and *j, σ*_*j*_ the fluid stress tensor, *p*_*j*_ the fluid pressure.

For *j* ∈ { *e, pa, pv*}, we assume that the transport of the solute is given by advection by the fluid and diffusion. Thus, we have

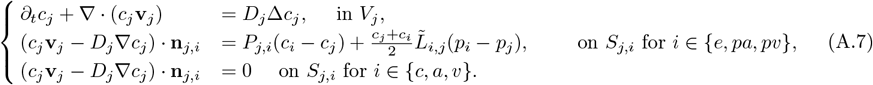

with *P*_*i,j*_ the diffusive permeability of the membrane and 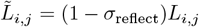 the convective permeability (with *σ*_reflect_ the solute reflection coefficient to represent the drag on the pores of the membrane that depends on the hydraulic radius of the solute).

To close the system, initial and boundary conditions must be given. However, to simplify the derivation, we will add them directly at the macroscopic scale.

To simplify the fluid problem, we assume CSF and interstitial fluid are incompressible, *i*.*e*. the micro-scale balance equation for the fluid reads

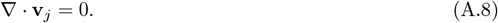

Furthermore, we assume that inertia is negligible so that momentum equation reduces to

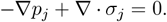

### A.3 Upscaling the microscopic problem

We first apply the volume averaging method (see Theorems 4-6) to the momentum equation to obtain

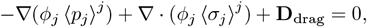

where we used 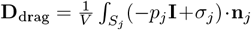 to account for shear stresses at the membranes separating the compartments or at the cells membrane (that do not absorb any fluid or solute). We further assume that these shear stresses can be represented as drag force (see *e*.*g*. [11])

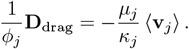

As pointed by Whitaker [67], we can assume that viscous forces ∇ · (*ϕ*⟨ *σ*_*j*_⟩ ^*j*^) are small compared to the drag forces when the length scale of the macroscopic system is large compared to the averaging volume taken into account, hence we arrive to

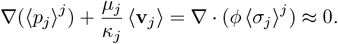

We obtain at the end the usual definition of the velocity field within a porous medium from Darcy [12]

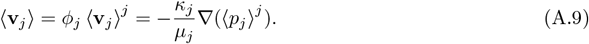

#### Remark 7

*This result is obtained for a non deformable medium as ϕ*_*j*_ *needs to be outside derivatives to be simplified*.

Applying volume averaging on Equation (A.8) and using the conditions at the interfaces *S*_*j,i*_, we have

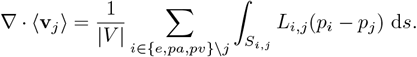

Assuming pressure is constant within each partial volumes *V*_*j*_ (but can vary from one volume to another), we simplify the previous equation to

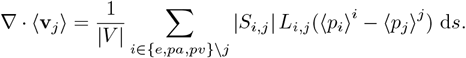

Denoting 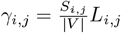 and using Equation (A.9), we arrive to

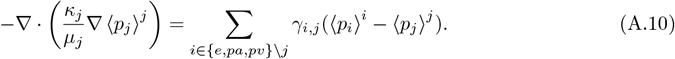

We now turn to the concentration equations. Applying volume averaging to System (A.7), we obtain for the time derivative

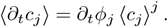

For the advection and diffusion terms, we have

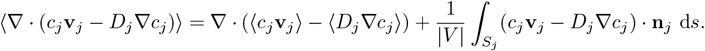

Assuming that the concentration is almost constant within *V*_*j*_ (*i*.*e*. we assume small microscopic fluctuations), the divergence term on the right-hand side of the previous equation simplifies to

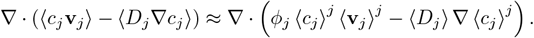

Furthermore, using the same assumption about small fluctuations within *V*_*j*_, and denoting 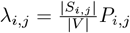 and 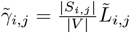, we have

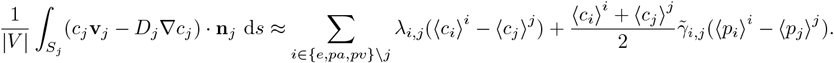

Altogether, we have for the macroscopic concentration problem

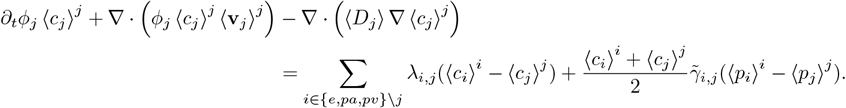

Denoting ⟨*D*_*j*_⟩ = *ϕ*_*j*_*D*_eff,*j*_ the effective diffusion tensor in compartment *j* as well as using Equation (A.9), we arrive to

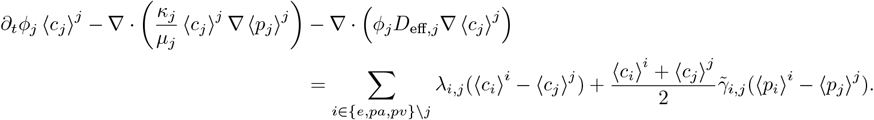

### A.4 The macroscopic model

Summarizing the derivation above, our macroscopic problem stated in terms of intrinsic averages reads for *j* ∈ {*e, pa, pv*}

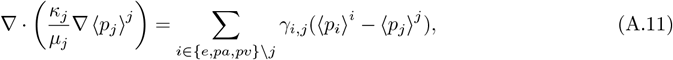

and

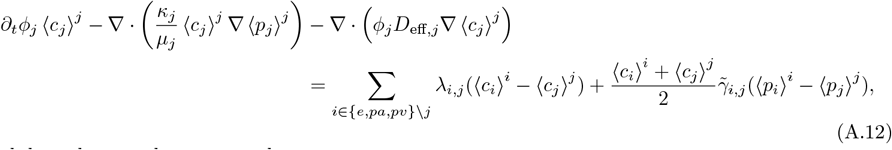

with boundary conditions given by

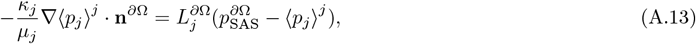

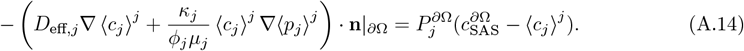

#### Remark 8

*As the boundary condition* (A.14) *gives the conditions for the intrinsic concentrations, we use in our model and scheme the boundary conditions for the macroscopic quantities*, i.e.

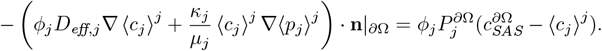

## B Details about the numerical scheme and our implementation

Our brain hemisphere mesh is obtained using the framework specified in [42] and the MRI data from the same book that is published at [41] under the Creative Commons Attribution 4.0 International license. To simulate the model (A.11)–(A.12), we employ the finite element method. To implement the scheme, we use the open-source finite element library Fenics [39, 1]. We use P1 finite elements for the pressure and concentration fields. The linear system associated to the finite element discretization of the pressure equations is solved using Conjugate Gradient with “hypre_amg” preconditionning. The linear system associated with the advection-diffusion part is solved using the GMRES algorithms with the ILU preconditioner. Our code is freely accessible at https://github.com/alexandrepoulain/glymphatics-glioma.

## C Parameter values for the three test cases

In the main body of this article, we modulated porosities and exchange parameters to represent the effect of the edemas and the tumor cells. We here provide tables summarizing the new parameter values for each test case.

For the pure nonvasogenic and the mixed edema test cases, we use the fluid and solute transfer coefficient between the ECS and vPVS in the nonvasogenic edema and the tumor regions indicated in Table 7, *i*.*e. γ*_*e,pv*_ = 7.51 × 10^−9^ and *λ*_*e,pv*_ = 3.11 × 10^−3^.

